# INSTRAL: Discordance-aware Phylogenetic Placement using Quartet Scores

**DOI:** 10.1101/432906

**Authors:** Maryam Rabiee, Siavash Mirarab

## Abstract

Phylogenomic analyses have increasingly adopted species tree reconstruction using methods that account for gene tree discordance using pipelines that require both human effort and computational resources. As the number of available genomes continues to increase, a new problem is facing researchers. Once more species become available, they have to repeat the whole process from the beginning because updating species trees is currently not possible. However, the *de novo* inference can be prohibitively costly in human effort or machine time. In this paper, we introduce INSTRAL, a method that extends ASTRAL to enable phylogenetic placement. INSTRAL is designed to place a new species on an existing species tree after sequences from the new species have already been added to gene trees; thus, INSTRAL is complementary to existing placement methods that update gene trees.

---

Methods for reconstructing species trees while accounting for gene tree discordance (Maddison, 1997) are now widely available (Szöllsi et al., 2014; Edwards et al., 2016; Warnow, 2017) and are adopted by many. Discordance-aware methods come in many forms, such as co-estimation of gene trees and species trees (e.g., Liu, 2008; Heled and Drummond, 2010; Boussau et al., 2013), site-based methods (e.g., Bryant et al., 2012; Chifman and Kubatko, 2014), and recently, direct modeling of polymorphisms (De Maio et al., 2013; Schrempf et al., 2016). The most scalable approach for species reconstruction has remained what has been called a summary approach (Mirarab et al., 2016): first gene trees are inferred independently for all loci, and then they are combined to build a species tree. Many methods are available for combining gene trees (e.g., Kubatko et al., 2009; Liu et al., 2009; Mossel and Roch, 2010; Liu et al., 2010; Chaudhary et al., 2010; Liu and Yu, 2011; Wu, 2012; Bayzid et al., 2013; Sayyari and Mirarab, 2016a), and many of them are statistically consistent under various models of genome evolution. In particular, many of the summary methods have established statistical guarantees (Liu et al., 2010; Allman et al., 2016; Shekhar et al., 2017) under the multi-species coalescent model (Pamilo and Nei, 1988; Rannala and Yang, 2003), which can generate incomplete lineage sorting (ILS) (Degnan and Rosenberg, 2009). Several summary methods, including ASTRAL (Mirarab et al., 2014b), NJst/ASTRID (Liu and Yu, 2011; Vachaspati and Warnow, 2015), and MP-EST (Liu et al., 2010) are in wide use.

Despite the progress for the *de novo* inference of species trees, updating trees has received little attention. As new genomes become available, researchers often need to know their position on an existing phylogeny. An expensive solution is to reconstruct the species tree from scratch each time new data becomes available. This process will require excessive computation and will not scale to groups with tens of thousands of genomes (more than a hundred thousand bacterial genomes are currently available).

A more efficient alternative is what has been called phylogenetic placement (Matsen et al., 2010): adding a new *query* species onto an existing phylogeny. For placing a new sequence onto a gene tree, we have maximum likelihood (ML) methods such as `pplacer` (Matsen et al., 2010) and `EPA` (Berger et al., 2011) and divide-and-conquer methods such as `SEPP` (Mirarab et al., 2012). Even earlier, sequential sequence insertion algorithms, which essentially solve the same computational problem, existed (e.g., Desper and Gascuel, 2002; Felsenstein, 1981). However, all these algorithms place a new sequence onto a gene tree. Adding a new species on a species tree is equally needed. We are not aware of any discordance-aware methods for placement onto species trees. Here, we present INSTRAL (Insertion of New Species using asTRAL) which extends ASTRAL to enable placing a new species onto an existing species tree.

## DESCRIPTION

### Background

ASTRAL estimates an unrooted species tree given a set of unrooted gene trees and is statistically consistent under the multi-species coalescent model given true gene trees (Mirarab et al., 2014b). ASTRAL seeks to maximize the quartet score: the total number of induced quartet trees in the gene trees that match the species tree. Similar to earlier work (Bryant and Steel, 2001), ASTRAL uses dynamic programming to solve this NP-Hard problem (Lafond and Scornavacca, 2017). However, to allow scalability, it constrains its search space so that the output draws its clusters from a predefined set *X*, which consists of clusters from gene trees and others that are heuristically selected (a cluster is one side of a bipartition). The most recent version, ASTRAL-III (Zhang et al., 2018) guarantees polynomial running time and scales to datasets with many thousands of species.

### Problem statement

*Quartet Placement Problem.* Given a set of *k* unrooted trees labeled with *n* + 1 species and a backbone tree on *n* species, find the tree that includes all *n* + 1 species and has the maximum quartet score with respect to the input trees.

Thus, one species, called the *query*, is not present in the backbone tree, and the goal is to insert the query species into the backbone. A typical use case of this problem is placing a new species onto an existing species tree (Fig. 1). Imagine a previous analysis has already produced a set of *k* gene trees on *n* species and an ASTRAL tree (inferred from those *k* gene trees). Now, a new species with genome-wide data has become available. To insert the new species onto a given ASTRAL tree, we first add it to each of the *k* gene trees using tools such as `SEPP`, `pplacer`, or `EPA`. Then, we use the updated gene trees in addition to the existing ASTRAL tree as input to the quartet placement problem; the output will be a species tree with the new species included. Just like ASTRAL, the use of the quartet score ensures that the inferred position of the new species is a statistically consistent estimator of its true position under the MSC model (given true gene trees).

**Fig. 1.**
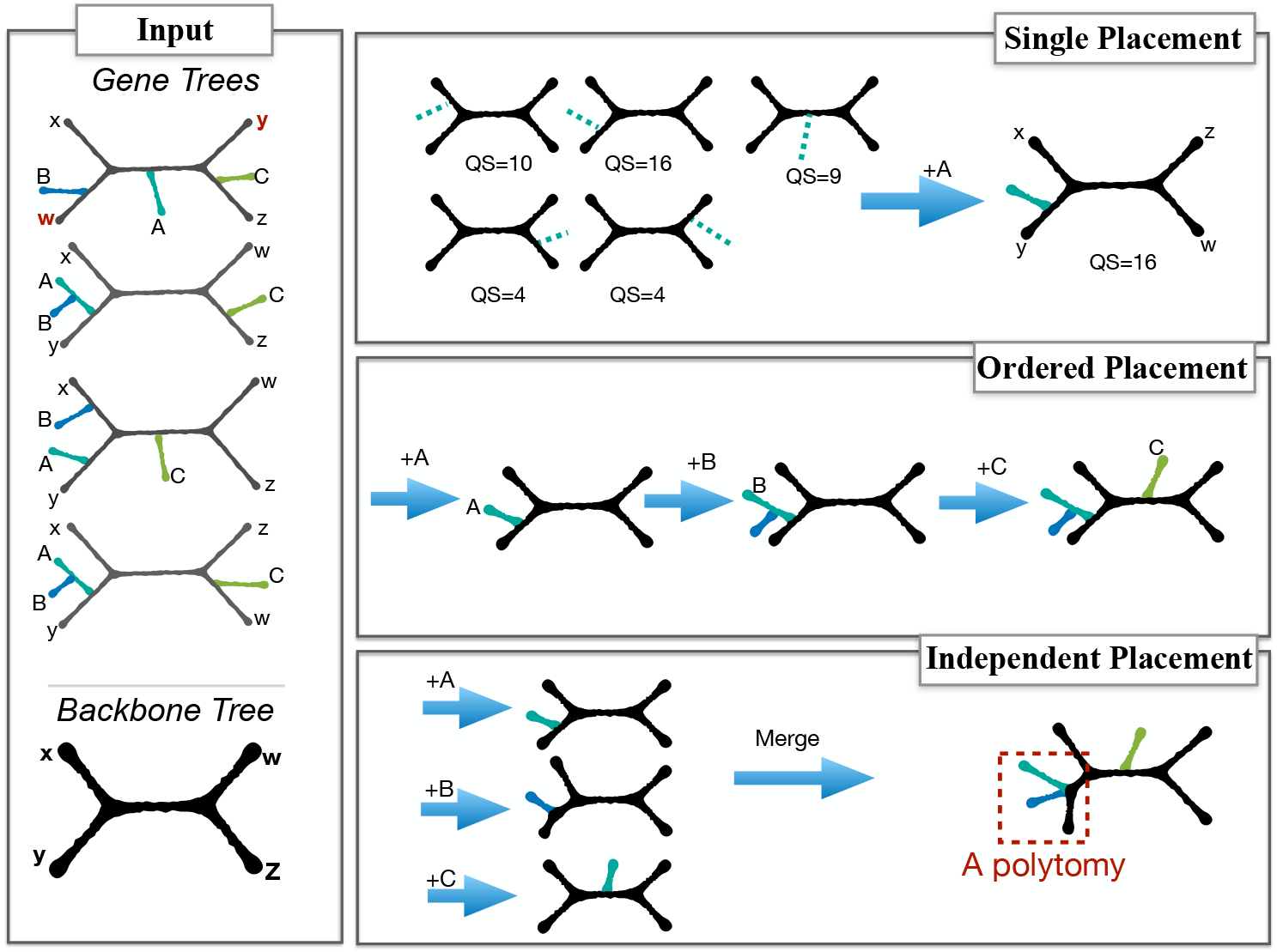
Left: The quartet placement problem. A backbone species tree with four leaves ({*x, y, w, z*}) and *k* = 4 gene trees are given; each gene tree also has new species (here, {*A, B, C*}). Note that the first gene tree is discordant with the species tree. Top right: placing a single new species (*A*) on the backbone tree requires computing the quartet score (QS) for each placement and finding the maximum. Here, the optimal placement is on the terminal branch of *y*, which matches 16 out of 20 quartets on {*x, y, w, z, A*} in the gene trees. Middle right: placing multiple species can be done by ordering them and placing them one at a time. Bottom right: alternatively, all new species can be placed independently, and the results can be merged at the end (creating polytomies when multiple new species are placed on the same branch).

### INSTRAL (single query)

INSTRAL finds the optimal solution to the quartet placement problem. Unlike AS-TRAL, the number of possible solutions to the placement problem is small (grows linearly with *n*), and thus, INSTRAL can solve the problem exactly even for large trees. In principle, it is possible to develop algorithms that compute the quartet score for all possible branches, one at a time, and to select the optimal solution at the end. However, the ASTRAL dynamic programming allows for a more straight-forward algorithm.

The ASTRAL algorithm will solve the placement problem if we define the search space (set *X*) such that *all* trees that induce the backbone tree and *only those trees* are allowed. To achieve this, *X* should be the set of all clusters in the backbone tree both with and without the new species added. More precisely, let *q* be a set with the new species and let *𝓑*(*T*) denote the set of all (including trivial) bipartitions of the backbone tree *T* on the leaf-set *ℒ* with each bipartition represented as a tuple: (*A, ℒ \ A*)*, A* ⊂ *ℒ*. Then

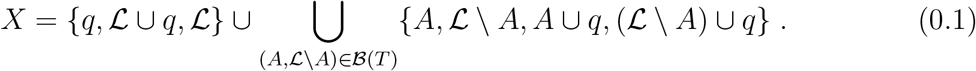

With this set *X*, the search space will include all possible placement of the query on the backbone tree (due to *A*∪*q* and (*ℒ\ A*) ∪*q*). Moreover, every bipartition built from *X* is one that existed in the backbone tree once *q* is removed and thus only trees that induce the backbone are allowed. Since ASTRAL finds the optimal placement restricted to the search space, this algorithm is guaranteed to solve the quartet placement problem exactly. The number of clusters in this search space is 3 + 4(2*n −* 3) = Θ(*n*). Thus, its running time increases as Θ(*n*.*D*) = *O*(*n*^2^*k*) where *D* is the sum of degrees of all *unique* nodes in the input gene trees (see Zhang et al., 2018, for details).

### Adding multiple new species

If multiple queries are available, we can still attempt to use the basic INSTRAL algorithm in one of two ways (Fig. 1). *i*) *Independent placement:* We add all the new queries independently without trying to find the relationship among the queries. This approach is reasonable if the goal is to detect the identity of some unknown species or if the set of new species are expected not to belong to the same branches of the backbone tree. If needed, we can merge separate placements into a single tree, introducing polytomies wherever multiple queries are placed on the same branch. *ii*) *Ordered placement:* We order the queries (e.g., arbitrarily) and then add them to the backbone one at a time, updating the backbone tree each time to include the latest query. This ordered placement approach gives us the relationships between queries. However, like similar greedy algorithms (Desper and Gascuel, 2002), it is not guaranteed to find the optimal tree at the end.

The advantage in using the independent insertion approach is that adding *m* queries requires time that increases linearly with *m* whereas the time needed for the ordered placement increases proportionally to *m*^3^. The *de novo* execution of ASTRAL-III on *n* + *m* species requires *O*(((*m* + *n*)*k*)^2.73^) time in the worst case (Zhang et al., 2018). In contrast, INSTRAL-independent would run in Θ(*m*.*D*.*n*) = *O*(*mn*^2^*k*) and INSTRAL-ordered would require *O*(*m*^3^*k* + (*n* + *m*)*nmk*). Thus, the relative running time of ASTRAL-III and INSTRAL-ordered depend on values of *n*, *m*, and *k*, while INSTRAL-independent is always faster than ASTRAL-III.

## BENCHMARK

### Datasets

We first benchmark INSTRAL on a simulated dataset previously generated by Mirarab and Warnow (2015). This dataset has 200 ingroup and an outgroup species and is generated using SimPhy (Mallo et al., 2015). By setting the maximum tree heights to 10^7^, 2 *×* 10^6^, or 5 *×* 10^5^ generations, this dataset has created three model conditions with respectively, moderate, high, or very high levels of ILS; the average normalized Robinson and Foulds (1981) distance (RF) between true gene trees and the true species tree are 15%, 34%, and 69%, respectively. In our experiments, we use gene trees inferred using FastTree-II (Price et al., 2010) from sequence data. These inferred trees have relatively high levels of gene tree error (25%, 31% and 47% for the three model conditions). We have 100 replicates per condition, and each replicate has 1000 gene trees, from which we have randomly sampled 200 and 50 gene trees to create three different input sets. Thus, in total, we have 9 model conditions (ILS level*×*# Gene). Following Mirarab and Warnow (2015), three replicates are removed because their gene trees are extremely unresolved; this leaves us with 9 *×* 100 *−* 3 *×* 3 = 891 datasets in total.

#### Leave-one-out experiments

For each dataset, an ASTRAL species tree inferred from gene trees is available. For each of the 200 ingroup species in each dataset, we prune it from the ASTRAL tree and by INSTRAL we add it back onto the tree, using FastTree gene trees as input. Thus, overall, we have 891 *×* 200 = 1.782 *×* 10^5^ independent placements. When there are multiple placements with equal quartet scores (happens in only 63 cases), we break ties similarly to the full backbone ASTRAL tree.

Among all of these placements, in only 316 cases (*<* 0.2%) the output trees have different quartet scores compared to the original ASTRAL tree. Note that INSTRAL is guaranteed to find the optimal placement, and theretofore, its quartet score is always at least as good as the ASTRAL tree. Thus, these 316 cases are those where ASTRAL has failed to find the optimal placement for a species. We note that 178 out of 316 cases correspond to the model condition with very high ILS and only 50 genes. Increasing the number of genes and reducing the amount of ILS both decrease the number of cases where ASTRAL is sub-optimal (Table 1). For example, with moderate ILS/1000 genes, only 4 out of 20,000 placements using INSTRAL improved quartet scores compared to ASTRAL. Only 176 of 316 cases result in any change in the RF distance of the inferred tree compared to the true tree, and only in 59 out of 176 cases did INSTRAL reduce the RF distance compared to ASTRAL. Thus, removing and reinserting a species using INSTRAL is generally consistent with the ASTRAL tree but in rare cases improves quartet scores.

**Table 1.**
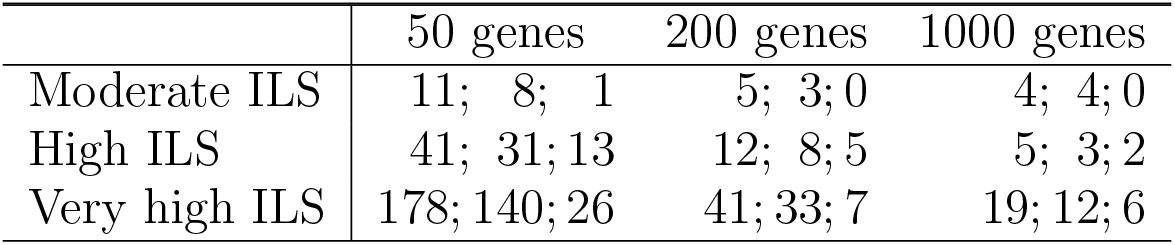
For each condition, we show the number of cases where (left) the INSTRAL tree has a different (i.e., higher) quartet score than the full ASTRAL tree, (middle) the Robinson and Foulds (1981) distance (RF) of the INSTRAL tree to the true tree is different than the RF distance of the full ASTRAL tree to the true tree, and (right) the INSTRAL tree has a *reduced* RF distance to the true tree compared to the full ASTRAL tree. All numbers are out of 20,000 insertions, except for very high ILS, which is out of 19,400.

|  | 50 genes | 200 genes | 1000 genes |
| --- | --- | --- | --- |
| Moderate ILS | 11; 8; 1 | 5; 3; 0 | 4; 4; 0 |
| High ILS | 41; 31; 13 | 12; 8; 5 | 5; 3; 2 |
| Very high ILS | 178; 140; 26 | 41; 33; 7 | 19; 12; 6 |

#### Ordered placement

To see if the agreement with ASTRAL remains if more species are placed using INSTRAL, we perform a second experiment. Here, we first prune a portion (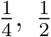, or 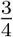) of species from the ASTRAL species tree, order removed species randomly, and then place them one after another on the backbone tree, updating the backbone tree each time (Ordered Placement in Fig. 1). In the end, we have a tree on the full leaf-set; this tree, which we call the INSTRAL tree, can be thought of as a greedy solution to the same problem ASTRAL seeks to solve.

ASTRAL and INSTRAL trees have similar RF distances to the true tree, but AS-TRAL is somewhat more accurate in the hardest conditions (Fig. 2a). Overall, the normalized RF error of ASTRAL is on average 0.3% lower than INSTRAL (corresponding to roughly half an edge), and these improvements are statistically significant (*p* ≪ 10^−6^ according to a paired t-test). Among all 891 × 3 = 2, 673 INSTRAL trees that we have computed, 1,470 have RF distances to the true tree that are identical to the ASTRAL tree. Differences in the RF distance are seen more often among replicates with very high ILS (mean RF difference: 0.7%), 50 genes (mean RF difference: 0.7%), or starting trees with 1*/*4 of the species (mean RF difference: 0.6%). Increasing the number of genes, increasing the size of the starting tree, and reducing the ILS reduce the number of mismatches between ASTRAL and INSTRAL (Fig. 2a).

**Fig. 2.**
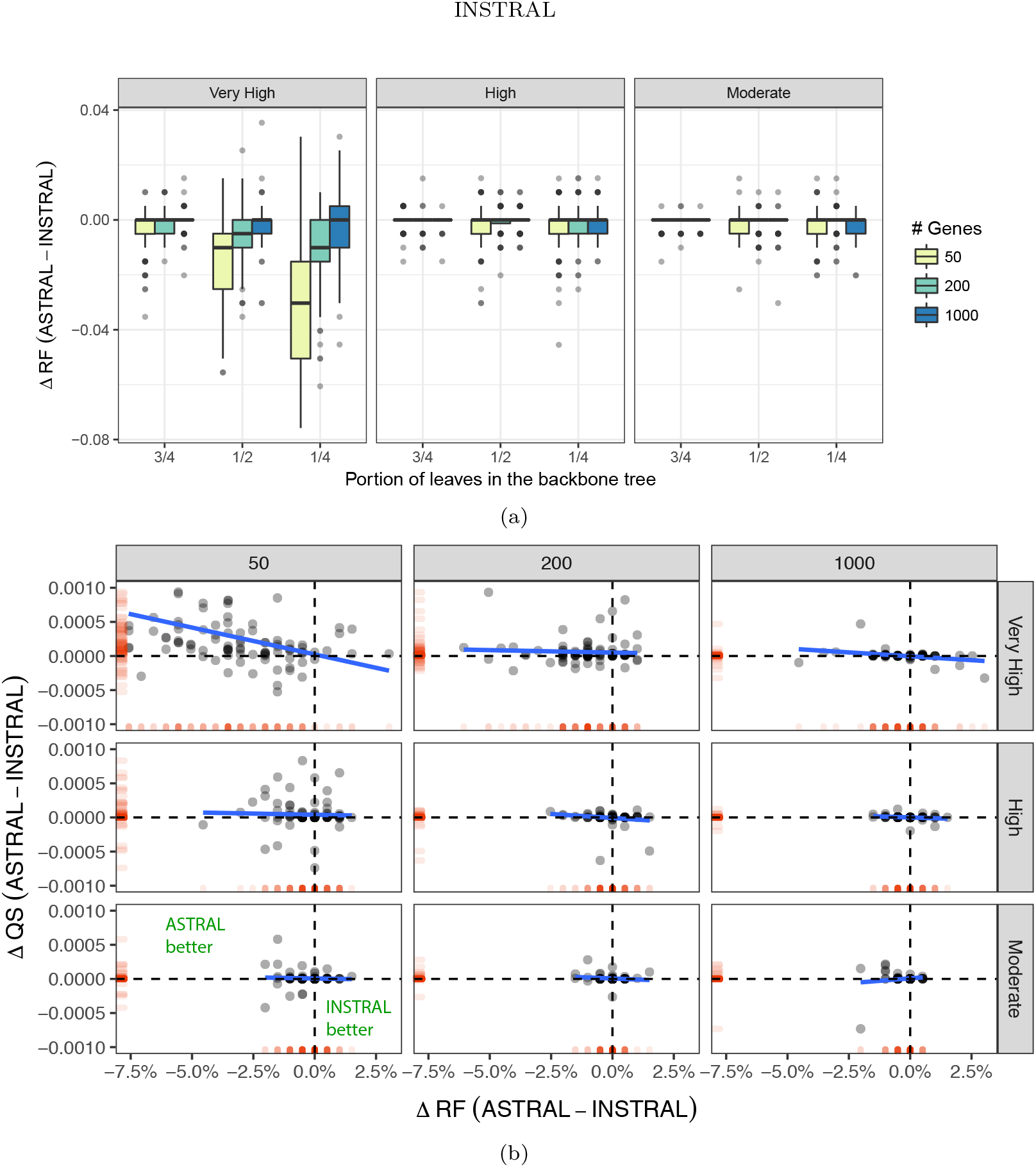
Comparison of ASTRAL and INSTRAL. (a) ∆ RF: The Robinson Foulds (RF) distance of the ASTRAL tree to the true tree minus the RF distance of INSTRAL-ordered tree to the true tree (negative: INSTRAL is better). The size of the starting tree is set to 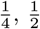, or 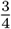 of species (51, 101, or 151). For three levels of ILS (boxes), each with three numbers of genes (colors), boxplots show distributions of ∆RF (100 points everywhere, except for very high ILS, where it is 97 points.) (b) Change in the quartet score (QS) versus the ∆RF for the starting tree with 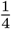 of species (see Fig. S1 for others). The marginal red bars show the projection of data on each axis.

Unlike the case of a single insertion, for multiple species, the quartet score of IN-STRAL can be higher or lower than ASTRAL. Overall, when the two trees do not agree, ASTRAL tends to have higher quartet scores (Figs. 2b and S1). Out of 2,673 cases, ASTRAL has higher quartet scores in 1,210 cases while INSTRAL is better in 231 (they tie in the remaining 1,232). Reducing the number of genes and increasing the level of ILS both magnify the improvements of ASTRAL compared to INSTRAL.

### Scalability

To test the scalability of INSTRAL, we started with a backbone tree of 10,000 species from a previous publication Zhang et al. (2018), and down-sampled it to smaller trees (down to 250). Each time, we placed 400 to 800 genomes on the backbone and computed the time INSTRAL took for the insertion (Fig. 3). On the backbone of 10,000 species, each placement took close to 16 minutes on average. As the backbone size decreased, the running time rapidly decreased and was close to 8 seconds on a backbone tree of 250 species. As expected, the running grows faster than linearly with the size of the backbone (proportional to *n*^1.3^ in this case).

**Fig. 3.**
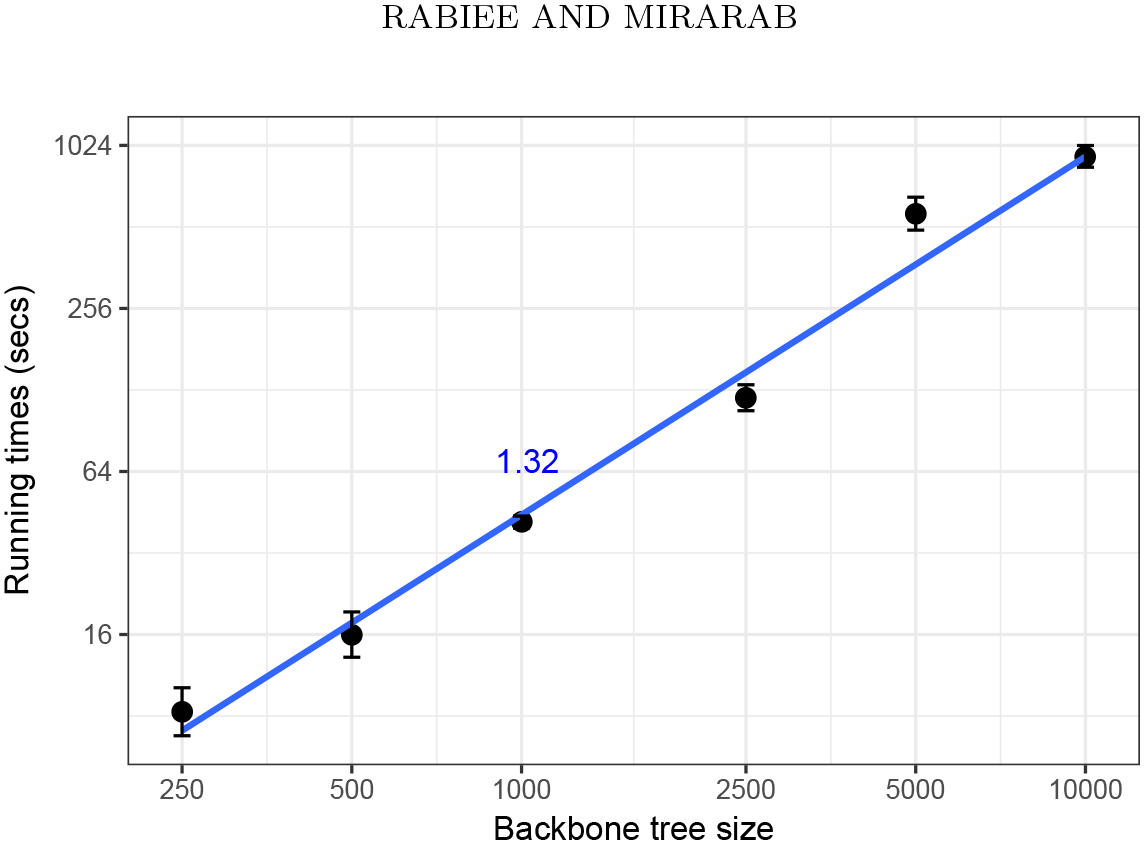
The running time scaling of INSTRAL versus backbone size *n*. Starting with a simulated dataset with 10,000 leaves, we prune random sets of leaves to create smaller trees. Dots and bars show the average and standard error of the running times of inserting a new genome to the backbone (800 insertions for *n <* 5000 and 400 insertions for n⩾ 5000). The slope (1.32) of the line fitted to this log-log plot gives an empirical estimate of the running time complexity being close to *n*^1.3^, which is consistent with the theoretical running time complexity of Θ(*n*.*D*) = *O*(*n*^2^).

## BIOLOGICAL EXAMPLES

We have tested INSTRAL on three biological datasets: two transcriptomic datasets on insects by Misof et al. (2014) and plants by Wickett et al. (2014), and an avian dataset by Jarvis et al. (2014). The insect dataset includes 1,478 protein-coding genes from 144 species spanning all of the insect diversity and has been recently re-analyzed using ASTRAL by Sayyari et al. (2017). The plant dataset includes 103 species and 424 genes, and the original study reported an ASTRAL tree. The avian dataset consists of 48 genomes representing all the orders of birds. For this dataset, statistical binning was used to build 2022 supergene trees (Mirarab et al., 2014a) and Sayyari and Mirarab (2016b) have published an ASTRAL tree on these supergene trees. Among these datasets, the avian dataset has extremely high levels of gene tree discordance.

For each of these datasets, we removed species one by one and placed them back onto the species tree using INSTRAL. In every case, INSTRAL found the same position for the new species as the backbone ASTRAL tree.

We also tested the ordered placement, where we randomly selected half of the species (20 replicates), removed them, ordered them, and inserted them back on the remaining part of the tree using INSTRAL. The resulting INSTRAL-ordered trees were similar to the full ASTRAL tree (Table 2), recovering the same tree in one-third of cases and changing by one or two branches in a majority of the remaining cases (Fig. S2a). In several replicates, trees changed for five or more branches, including two replicates of the avian dataset, where the INSTRAL differed from ASTRAL in nine branches. In both cases, two or three unstable taxa had moved by several branches, causing the high incongruence (Fig. S2b). More broadly, changes are mostly among unstable taxa. For example, in the avian tree, Hoatzin, the most challenging taxon, moves by one branch in several replicates. The resulting INSTRAL trees have reduced quartet scores compared to ASTRAL trees (Table 2). Overall, these results indicate that for datasets with very high ILS, using INSTRAL instead of ASTRAL runs the risk of producing sub-optimal trees.

**Table 2.** The average and standard deviation of RF distance between ASTRAL and INSTRAL trees as well as the change in the quartet score (INSTRAL-ASTRAL) on 20 random sets from each biological dataset. For each random set of leaves as a backbone tree, ordered placement has been done.

|  | Avian |  | Insects |  | 1KP |  |
| --- | --- | --- | --- | --- | --- | --- |
|  | mean | stdev | mean | stdev | mean | stdev |
| RF | 0.0311 | 0.0592 | 0.0195 | 0.0126 | 0.0115 | 0.0131 |
| QS | -0.0001098 | 0.0002973 | -0.0000026 | 0.0000361 | -0.0000039 | 0.0000057 |

## AVAILABILITY

INSTRAL is open-source and freely available on GitHub (https://github.com/maryamrabiee/INSTRAL). It is implemented in Java with straight-forward installation (the only dependency is Java 6+). A template tutorial and instructions to run INSTRAL is given there. The generated data, scripts to generate those data and results given in this paper are also available on GitHub (https://github.com/maryamrabiee/INSTRAL-results).

### Acknowledgments

This work was supported by the NSF grant IIS-1565862 to SM and MR. Computations were performed on the San Diego Supercomputer Center (SDSC) through XSEDE allocations, which is supported by the NSF grant ACI-1053575.

---

**Fig. S1.**
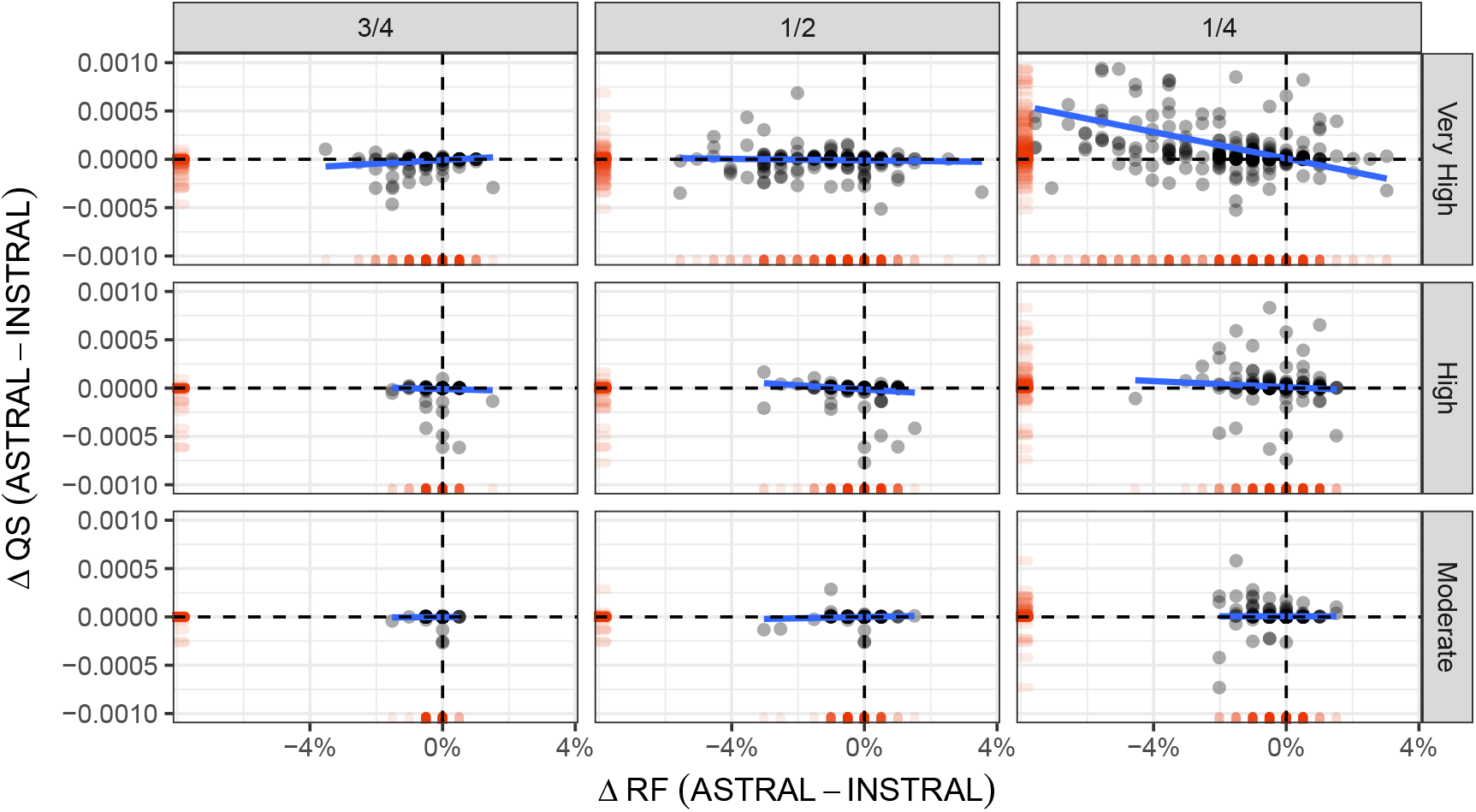
∆*QS* of the output trees of the two methods based on ∆*RF* in all model conditions and with different backbone trees containing 1*/*4, 1*/*2 and 3*/*4 of the species. The red tone on the x and y axes is the projection of data on both axes and represents the distribution of data on each axis.

**Fig. S2.**
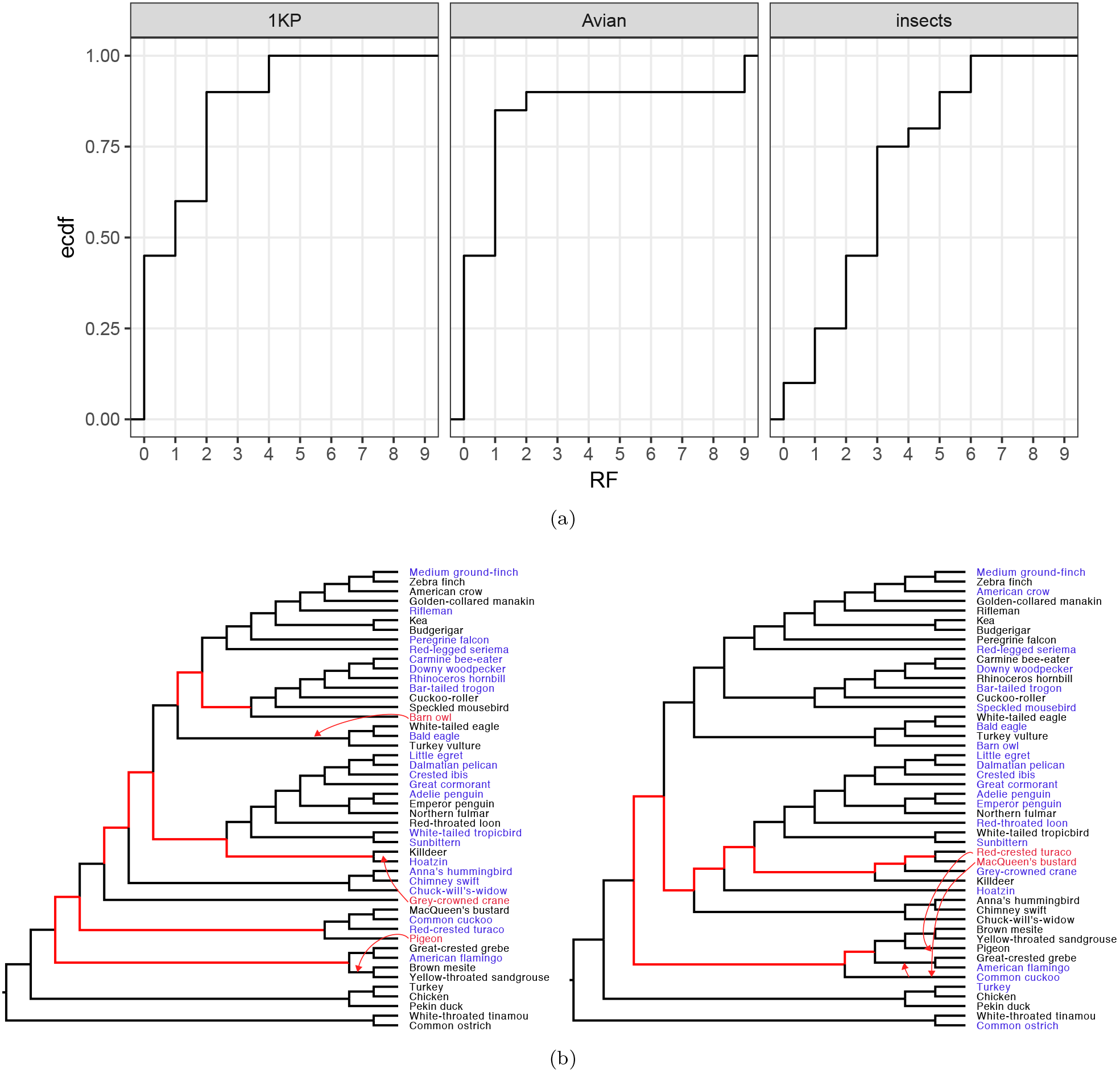
Results on the biological dataset. (a) The empirical cumulative distribution function (ecdf) of the number of branch differences between running ASTRAL and INSTRAL-ordered (x-axis) on the three biological datasets. The distributions are over 20 replicates, each using half of the species, selected randomly, as backbone and placing the rest in some random order. (b) Two INSTRAL avian trees had high levels of difference from ASTRAL. Blue: backbone species. Red edges: different from ASTRAL. Red tips: The entire differences can be described by three (top) or two (bottom) rouge taxa that have moved far away from their position in the ASTRAL tree (red arrows).

